# Forensic post-mortem interval (PMI) estimates: variation in fly developmental times of individuals

**DOI:** 10.1101/064782

**Authors:** Mihály Földvári

## Abstract

Immature stages of flies are paramount in establishing the post-mortem interval (PMI) in forensic practice. My focus is on differences in developmental time that can be influenced by genetic differences or individual life history traits, which latter may be interpreted as life history decisions.

Data of a calliphorid fly species (Lucilia ampullacea) are presented: one female produced 300 eggs within an hour and the individual developmental time varied subsequently to a great extent – when the first flies emerged from their puparia there were still first instar larvae in the food (pig liver) provided.

In conclusion the estimated PMI must be based on a wide range of collected flies (not simply the oldest or largest or widest individual), since a limited sample can be one extremity of a potentially bell shaped (Gaussian) frequency distribution of developmental times — unrepresentative sampling will bias the PMI in an unpredictable way. One possible solution can be to use large, randomized samples and their body measurement means.

## INTRODUCTION

In forensic entomology a major subject is the accurate estimation of PMI (post-mortem interval), the time between death and the discovery of a corpse, and this can be influenced by several modifying circumstances (Catts, 1992). In case of immature insects the PMI is further divided into the pre-appearance interval and the development interval (Matuszewski, 2011).

Among the factors to be considered one has not been treated in appropriate detail: the variation in the amount of time required for the dipterous larvae to develop into pupariums.

One general method in PMI estimation is to collect larval samples at the time of discovery and then calculate the possible time of death or exposure of the corpse to oviposition (uncovering or relocating the corpse). As a general rule usually larger (older) maggots are used as indicators of PMI, since they are considered to require more time for development (Catts, 1992; Villet et al, 2010; Rivers and Dahlem, 2014).

One major influence on the calculations can be that there is evidence for precocious larval development, which means that the PMI can be overestimated because of those larvae that started development before oviposition (Wells and King, 2001).

The uncertainty of the PMI estimate can be further increased by the variation in individual larval development lengths: Here I show an example of *Lucilia ampullacea* from the family Calliphoridae within the order of Diptera. It is not the most common fly to be found on corpses placed outdoors, but they are not rare under such circumstances, especially in Central Europe (Szpila, 2010).

## Materials and Methods

The eggs have been collected from one female *Lucilia ampullacea* found in the Botanical Garden of the University of Debrecen (Debrecen, Hungary) on 24^th^ October 2013. The fly was transferred to a plastic laboratory cage (25°C and 45-55% humidity, 12h : 12h day : night photoperiod) with the dimensions 45cm × 20cm × 20cm, where it laid eggs in one hour and then died. The substrate was pig liver for the egg laying and for the larval development with sufficient quantity to serve as food source for all of the larvae.

The developmental stages and numbers were observed and counted during the following days to establish developmental times and frequency distribution of times required to reach different stages (puparium and adult fly).

The number of eggs has been established by counting them in groups of 10 (they were not relocated), therefore it is not accurate, but a very good estimate. The counting of pupariums occurred on every day in the morning, and these pupariums have been removed from the substrate to avoid confusion about the numbers each day, but they were kept within the plastic laboratory cage.

## Results

Out of the 330 eggs laid there were 264 pupariums formed, many of which developed into adult flies (not counted). The frequency distribution of the puparium numbers on each day is shown in Fig. 1. The first flies emerged from the pupariums on the 8^th^ day after the eggs were laid.

**Figure 1.**
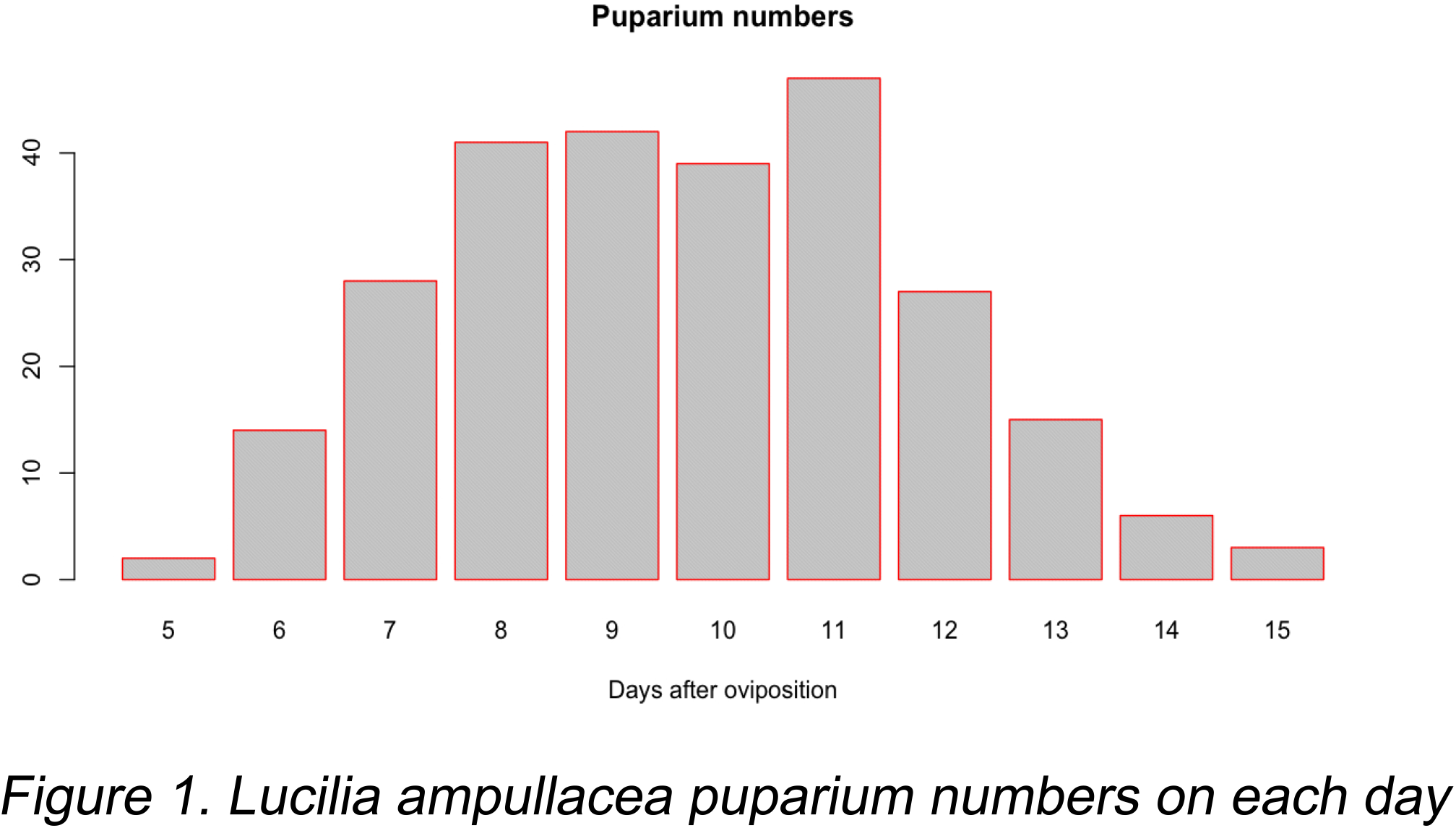
Lucilia ampullacea puparium numbers on each day.

## Discussion

Even though the fly eggs were laid within one hour, the development of the immature larvae (until puparium) took 5 to 15 days for the individuals. This is a distinct difference, since the range of the developmental age of the individuals increased from one hour to nine days from the first egg laid to the formation of the last puparium.

This is a very important observation in terms of PMI estimation, since data based on larval or puparium numbers at a particular time may not be accurate if the sample size is too low.

It is not an exact amount of time a larva needs for development, but a distribution of these times with a minimum, maximum and an average time in a shape of a frequency curve (e.g. Fig. 1.). In order to establish a correct age for the larval population on a potential corpse, there have to be enough flies sampled, so that the distribution curve can be produced, and then the approximate time of oviposition can be estimated.

During laboratory tests of larval development several studies use techniques that reduce the variance by not feeding the females prior to oviposition (Brown et al., 2015) or selecting only 1h old larvae (Richards et al., 2009). These circumstances cannot be reproduced under natural conditions and laboratory measurements will necessarily differ from the natural ones, since they reduce the above-mentioned variance of the developmental times.

The general usage of the oldest (largest) larva (Catts, 1992; Villet et al, 2010; Rivers and Dahlem, 2014) can be misleading, since it may not represent the oldest individual on the scene, just as likely as a smaller maggot may have spent the same amount of time there: because of the developmental variation they both may have been there for that long, but have reached a different stage. Davies and Ratcliffe (1994) report on this potentially large variation, but they only suggest using the upper quartile (body weight) of a large sample to reduce these effects, and do not go into details of the frequency distribution curve.

The best approach to solve this problem can be running tests for the standard distribution curve of developmental times to be established and then the measured case data from a large sample collected from the corpse can be compared to the standard.

The level of uncertainty given for the PMI estimate has to be adjusted with the variance introduced by the frequency distribution of the larval sample. This is further complicated with multiple egg laying events by the females on a corpse. The flies continue to lay eggs, and the developmental times of these occurrences will overlay on each other. Each oviposition event will have its own range and variance, laid on top of the other (not to mention that multiple species will produce eggs on the same corpse).

If the sample size is large enough and the individuals are randomly selected from the source, these problems can be minimised.

